# Disentangling choice value and choice conflict in sequential decisions under risk

**DOI:** 10.1101/2021.10.13.464217

**Authors:** Laura Fontanesi, Amitai Shenhav, Sebastian Gluth

**Affiliations:** Department of Psychology, University of Basel; Department of Cognitive, Linguistic, and Psychological Sciences, Brown University; Department of Psychology, University of Hamburg

**Author notes:** shared senior authorship (Amitai Shenhav, Sebastian Gluth). Amitai Shenhav acknowledges support from the NIMH (Grant R01MH124849) and from an Alfred P. Sloan Foundation Research Fellowship in Neuroscience. Sebastian Gluth acknowledges support from the European Research Council (ERC) under the European Union’s Horizon 2020 research and innovation program (Grant agreement No. 948545).

**Keywords:** foraging, decision difficulty, choice conflict, response times, risk, sequential sampling models

## Abstract

Recent years have witnessed a surge of interest in understanding the neural and cognitive dynamics that drive sequential decision making in general and foraging behavior in particular. Due to the intrinsic properties of most sequential decision-making paradigms, however, previous research in this area has suffered from the difficulty to disentangle properties of the decision related to (a) the value of switching to a new patch versus (b) the conflict experienced between choosing to stay or leave. Here, we show how the same problems arise in studies of sequential decision-making under risk, and how they can be overcome, taking as a specific example recent research on the ‘pig’ dice game. In each round of the ‘pig’ dice game, people roll a die and accumulate rewards until they either decide to proceed to the next round or lose all rewards. By combining simulation-based dissections of the task structure with two experiments, we show how an extension of the standard paradigm, together with cognitive modeling of decision-making processes, disentangles value-from conflict-related choice properties. Our study elucidates the cognitive mechanisms of sequential decision making and underscores the importance of avoiding potential pitfalls of paradigms that are commonly used in this research area.

## Introduction

Sequential decision making refers to situations in which we continue to take a series of similar actions until we either decide to stop or are required to do so. A very prominent case of sequential decision making is foraging. A foraging animal collects food from their current patch of land (e.g., a grazing area), where resources are gradually depleting, until it decides to leave this patch for a new one, or until no resources are left. Humans face similar dilemmas on a regular basis. A characteristic example are housing bubbles. The longer we wait to sell a house, the higher the potential gain as long as the prices keep increasing. But if the housing market collapses, we might have to sell the house at a price that is even lower than the acquisition costs. Critically, both the grazing animal and the house seller need to trade off the expected benefits and risks of staying in the current situation against those of making a switch (Charnov, 1976). An inherent property of sequential decisionmaking problems is that this trade-off between staying and switching becomes increasingly difficult the longer one stays in a given environment (Shenhav, Straccia, Cohen, & Botvinick, 2014). Thus, when the animal starts to exploit a new and rich patch, it is obvious that the animal should keep harvesting from the patch for a while. However, as the patch gets more and more depleted, the expected benefits from staying and switching become increasingly similar. The *marginal value theorem* predicts that a reward-maximizing animal will leave the old patch as soon as the expected benefits of switching is equal to or higher than the expected benefits of staying (Charnov, 1976). As a consequence, the final decision is usually made at the point of maximal choice conflict, that is, at the indifference point (IP) between staying and switching (Shenhav et al., 2014).

This feature of sequential decision making has been the center of a recent controversy on the neural basis of foraging (Kolling et al., 2016; Shenhav, Cohen, & Botvinick, 2016). In particular, it has been debated whether the dorsal anterior cingulate cortex (dACC) encodes either the subjective value of switching to the next patch (henceforth: ‘switch value’) (Kolling, Behrens, Mars, & Rushworth, 2012) or the conflict of choosing whether to stay or to switch (Shenhav, Straccia, Botvinick, & Cohen, 2016; Shenhav et al., 2014). The methodological challenge behind this debate is that both the switch value and the conflict rise monotonically until the IP, and thus can be confounded when used as predictors for neural activity (Figure 1). The two can be dissociated, however, by modifying standard foraging tasks so that more decisions are made *after* the IP. This is because after the IP, the switch value keeps increasing while conflict decreases (as it becomes more and more obvious that switching is the preferable choice). Thus, switch value and conflict become deconfounded.

**Figure 1.**
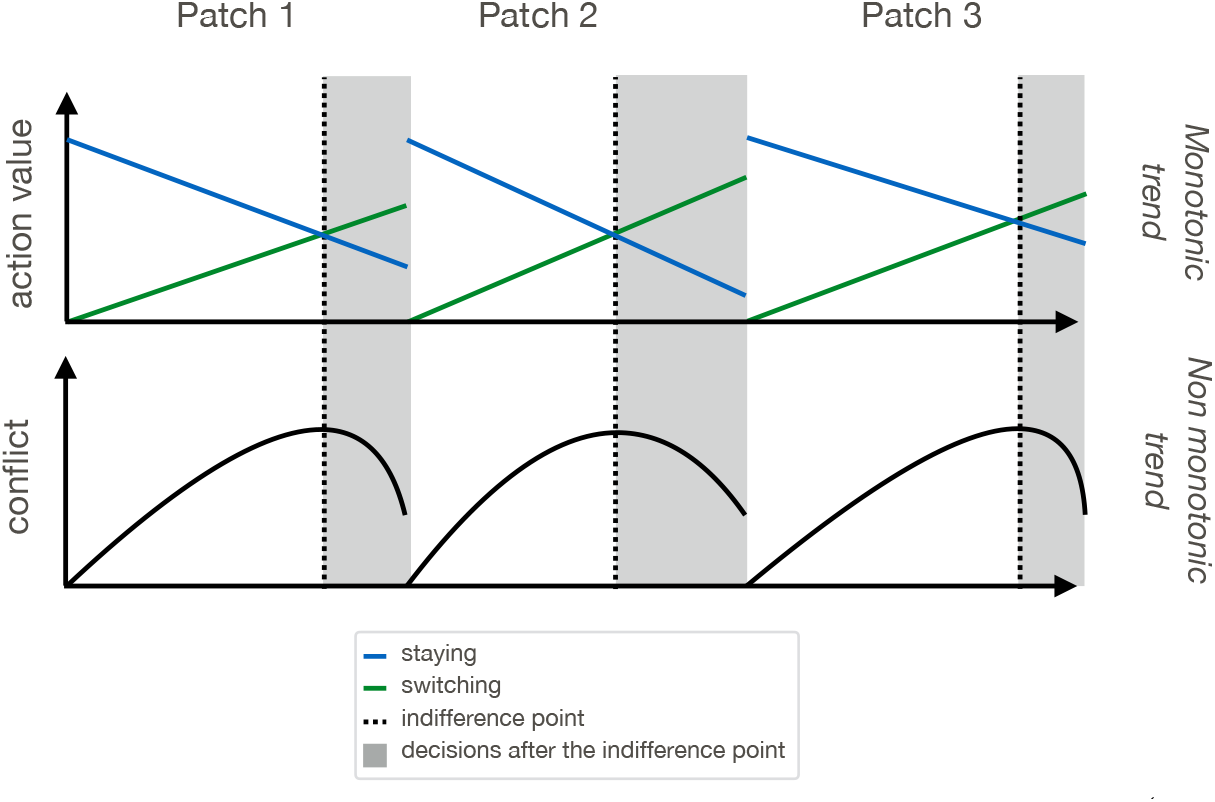
Shaded grey areas represent decision settings after the IP (i.e., the point at which leaving the patch has the same expected utility as staying in the patch). Decisions made after the IP have lower conflict than those made at the IP, and are thus crucial to discriminate the monotonically increasing switch value from the non-monotonic effect of choice conflict.

Notably, the same methodological challenge can also be found in sequential decision making under risk and uncertainty. A prominent example of this type of task is the Balloon Analogue Risk Task (BART), which involves inflating a virtual balloon with sequential pumps. Participants accumulate more money the larger the balloon gets, but lose that accumulated money if the balloon pops (Lejuez et al., 2002). Thus, each decision to pump the current balloon rather than cash in and move on to the next balloon comes with potential risk and reward. Consistent with the foraging literature discussed above, neural signals that increase with every pump have been found in dACC (Schonberg et al., 2012), but it remains unclear whether these signals represent the value of cashing in (i.e., switch value), choice conflict, escalating risks, or simply the number of executed pumps per round.

A recent study (Meder et al., 2016) attempted to tackle this question using a similar task, based on the dice game ‘pig’. In this game, participants repeatedly roll a die in a series of rounds. Every time they roll a die and a number between 2 and 6 faces up, that number is added to their accumulated reward for that round. If they roll a 1, however, the round ends and they lose the rewards accumulated in that round. After every roll of a die that does not result in a 1, participants can choose whether to terminate the current round or to continue rolling the die (Figure 2A). If they decide to terminate, the cumulative sum of rewards is cashed in. Their ultimate payoff depends on the average cumulative sum of rewards per round, so there are costs both to rolling a 1 (and thus losing everything) as well as cashing in too early (and thus collecting too little for that round). In fact, the expected value (EV) maximizing policy for a risk-neutral player is to keep rolling until the expected loss amount is higher than the EV of one more roll (Meder et al., 2016). Thus, the ‘pig’ game exhibits a simpler and less ambiguous definition of the optimal policy compared to the BART. Still, it shares the BART’s sequential decision-making structure (and that of foraging tasks), and with it the difficulty to separate choice conflict from switch value. Regardless, Meder and colleagues (2016) reported that neural activity in dACC scaled linearly with the cumulative sum of rewards in each round (which effectively represents the switch value in this task), and found a significant correlation between the cumulative sum of rewards and response times (RTs). They concluded that their data provided evidence in support of the switch-value hypothesis and against the conflict hypothesis (Kolling et al., 2012).

**Figure 2.**
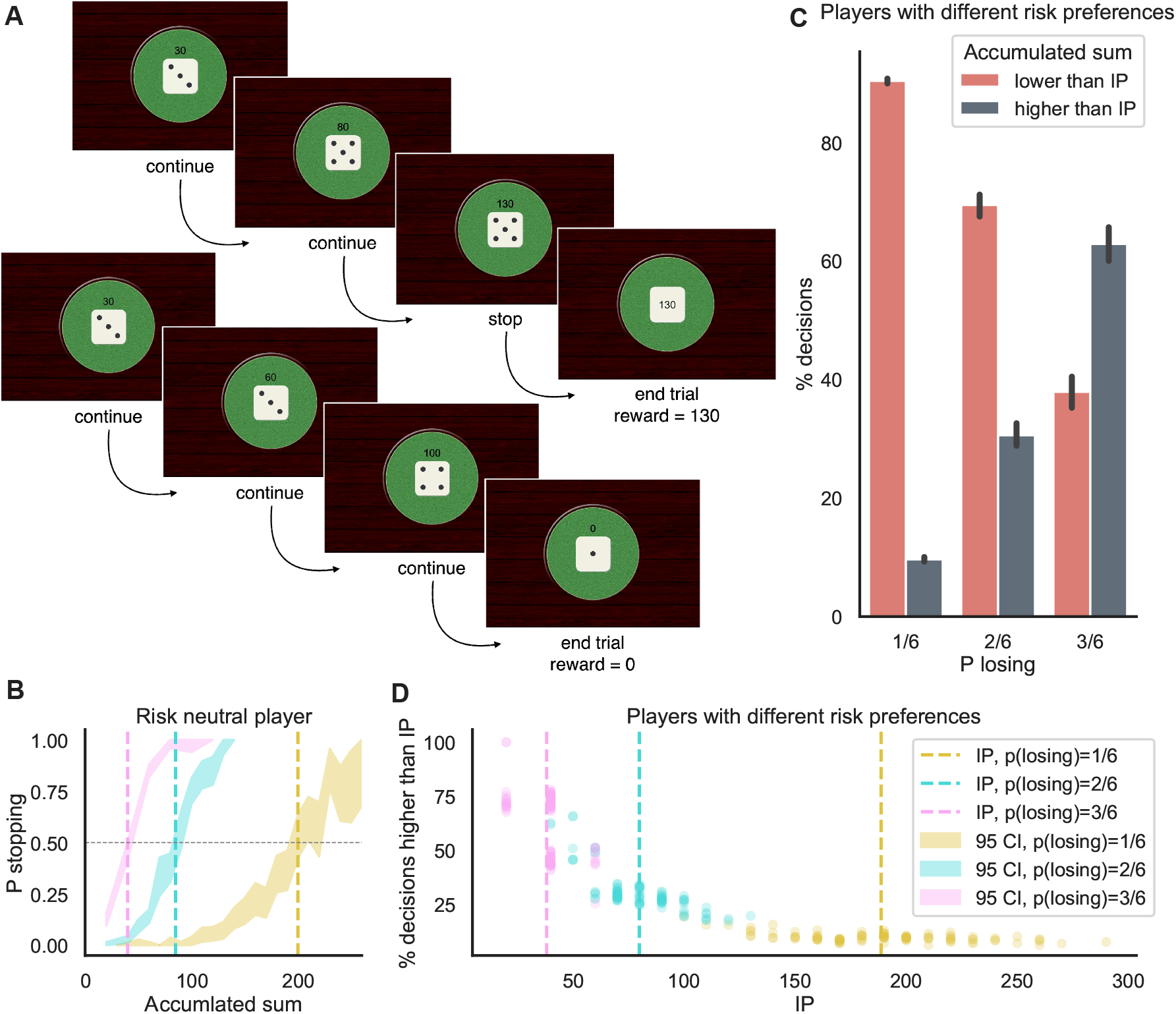
**A**: Example of two rounds of the ‘pig’ dice game, the top one ending with a decision to stop and a reward of 130, the bottom one ending because of a rolled 1 and with no reward. **B** Probability of stopping (shaded areas) as a function of the cumulative sum within a round for a simulated risk-neutral player. The IPs (dashed lines) for the 1/6, 2/6, and 3/6 conditions are, respectively: 40, 85, and 200. **C**: Percentage of decisions before and after the IP for a simulated group of players with different risk preferences (risk neutral on average). The number of decisions made after reaching the IP increases with the probability of losing, making the task more balanced. **D**: Percentage of decisions made after the IP as a function of the IP itself, for the same group of simulated participants as in C. Participants with higher IPs (more risk seeking) experience a more imbalanced task in the 2/6 and 1/6 conditions.

The purpose of the current study was to stress the need for careful consideration of a task’s structure as well as to demonstrate the benefits of employing computational modeling of behavior when studying sequential decision making. To this end, we used the ‘pig’ dice game as an exemplary sequential decision-making paradigm and subjected it to careful theoretical and empirical examination. In the following, we will first show that the standard version of the ‘pig’ dice game cannot dissociate variability in switch value (or in the number of choices to continue within a given round) from variability in choice conflict. As a consequence, it is very difficult to disentangle the relative contributions of these variables to neural activity measured while performing the task, as in Meder et al. (2016). Second, we will introduce a simple extension of the paradigm to mitigate this shortcoming. Third, we will present results from two behavioral experiments, one a replication of the original and the other a novel extension of the task that was designed to better discriminate switch value and conflict. Finally, we will use modeling of choice and RT data via the diffusion decision model (DDM, Ratcliff (1978); Ratcliff and Rouder (1998)) to map these different task dynamics onto parameters of a well-established computational model of decision making.

## Results

### Structure of the ‘pig’ dice game and modifications

In the version of the ‘pig’ dice game used by Meder and colleagues, the dice numbers were multiplied by 10 to specify the rewards. For example, cashing in after rolling a dice three times with numbers 3, 5, 5 would lead to a cumulative sum of rewards of 30+50+50 = 130 (top example in Figure 2A). On the other hand, if a 1 is rolled after 3, 3, 4 are rolled, the reward is not 110 but 0 (bottom example in Figure 2A). The time that participants spent on the entire task was fixed, and the monetary payoff was based on the average number of rewards collected per round. This combination of task design and incentive structure implies a fairly simple definition of the optimal strategy for a risk-neutral player who seeks to maximize EV. The player should continue rolling the die until a cumulative sum of rewards of 200 has been reached and then terminate the current round by cashing in (i.e., IP = 200). The rationale behind this EV-maximizing strategy is that the expected loss of rolling a 1 on the next try exceeds the expected gain of rolling one of the other numbers as soon as more than 200 points have been accumulated.

In this standard version of the game, it takes 4-10 consecutive die rolls to accumulate 200 points. Once that number is reached or crossed, points accumulate at a similarly gradual pace. Given that subjects are incentivized to avoid too many die rolls (because additional die rolls carry the risk of rolling a 1), and given this gradual pace of point accumulation, it is unlikely that a participant will continue to accumulate a substantial number of points past their IP. In other words, the design of this task limits the number of decisions that can be observed after the IP. This presents a significant challenge for disentangling correlates of reward accumulation and choice conflict, because it means that having greater accumulated reward will also generally mean being nearer to one’s IP (i.e., in a state of greater choice conflict about whether to continue or stop).

To demonstrate this, and to quantify the relationship between reward accumulation and choice conflict under different task conditions, we simulated a participant playing this game who chooses to continue or to stop in a probabilistic manner depending on the current distance of the cumulative rewards from 200 (the risk-neutral IP, details provided in Methods, Figure 2C, yellow curve). On average, this participant made only 9.7% of their decisions to stop the round past this IP. We then varied these risk preferences (IPs) across 100 simulated players and found that, across this population, players made only between 6% and 19% (7% on average) of their decisions past their IP (Figure 2C, where P losing = 1/6). This low proportion of choices past IP was higher when for risk-averse participants (i.e., IP < 200) compared to risk-seeking participants (i.e., IP > 200) (see Figure 2D, yellow dots).

To mitigate the imbalance of decisions before and after the IP, one can simply increase the probability of losing the cumulative sum of rewards at each roll of the die. In the standard version, participants have a 1/6 chance of losing with each roll (i.e., if they roll a 1). If this probability is increased to 2/6 (e.g., lose with a roll of 1 or 3) or to 3/6 (e.g., lose with 1, 3, or 5), then the optimal strategy is to shift one’s IP substantially downward. For a risk-neutral participant, this IP would be 85 points for the 2/6 case or 40 for the 3/6 case (Figure 2B, pink and blue curves). As a consequence, participants are expected to make fewer decisions prior to reaching their IP (because these lower IPs will be reached with fewer die rolls) and thus to have more balanced numbers of decisions before and after reaching their IP. Indeed, our simulations predict that participants will make 30.6% of their decisions past their IP when the probability of losing is 2/6, and that this number increases to 62.8% when the probability of losing is 3/6 (Figure 2C and D).

We tested these predictions empirically across two experiments. Experiment 1 was a replication of the study by Meder and colleagues. The purpose of this experiment was to test whether we can replicate the central results of the original study. We also sought to substantiate our critique of the standard version of the task with empirical data. In Experiment 2, we tested the standard version together with the two proposed changes of loss probabilities as described above.

### Experiment 1: replication of the standard ‘pig’ dice game

Participants (N=30) performed the standard version of the ‘pig’ dice game with a constant 1/6 probability of losing with each roll. To estimate each participant’s IP, we regressed the decision to stop or to continue onto the cumulative sum of rewards for a given round, using a hierarchical Bayesian logistic regression model (details provided in Methods). The cumulative sum of rewards was a good predictor for the probability of stopping, as indicated by a well-above-zero regression coefficient (HDI = [0.03, 0.05], Figure 3A). In line with Meder et al. (2016), we found that the average estimated IP was below the EV-maximizing IP of 200 (HDI = [92.73, 163.06], Figure 3B), indicating risk aversion. In line with our simulations, there were only very few decisions made after the IP (M=17.1%, SD = 10.7%, Figure 3C).

**Figure 3.**
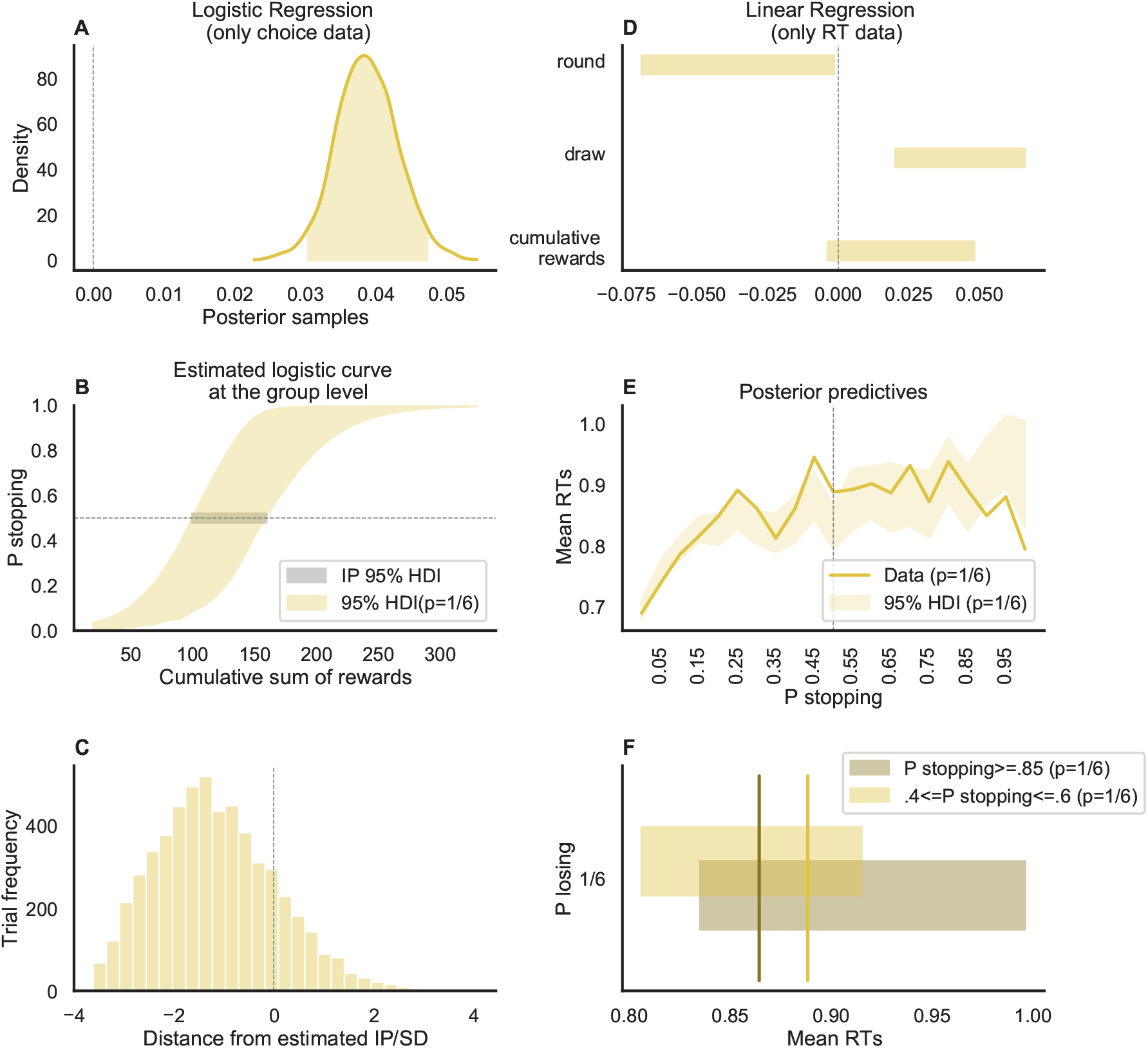
Regression analyses Experiment 1. The left column shows the results of the logistic model fit on choice data, while the right column shows the results of the linear model fit on RTs. **A** Posterior distribution of the cumulative sum of rewards coefficient at the group level (the shaded area is the 95% HDI), when we predict the probability of stopping within a round. **B** Estimated logistic curve (colored shaded area) and the IP (grey shaded area) at the group level. **C** Distribution of trials before and after the estimated IP. **D** Posterior distributions of the round number, draw number within a round, and cumulative sum of rewards coefficients at the group level (the shaded area is the 95% HDI), when we predict RTs. **E** Posterior predictives of mean RT data as the probability of stopping increases within a round, against the mean RT data. **F** Comparison of the same posterior predictives, selectively at the points of maximum conflict and at the points of maximum probability of stopping. Here the vertical lines represent the data while the shaded bars are the predictions.

The low number of decisions made after reaching the IP confirms our simulation results reported above and call into question whether the standard version of the task is suitable to isolate the influence of choice conflict from other factors that drive behavior, such as the expected utility of stopping. Nevertheless, we tried to identify such potential effects by analyzing the RT data of Experiment 1. More specifically, we fitted a hierarchical Bayesian linear regression model that regressed log(RT) onto the predictor variables round number, within-round draw number, and cumulative sum of rewards. By doing so, we found influences on RT for all predictor variables except for the cumulative sum of rewards (see Figure 3D and Table B2). A careful look at the development of RT over the course of the round of the game indicates that it is difficult to decipher whether RT decreased beyond the IP (Figure 3E), as would be expected given the lower number of trials after the IP. More specifically, when comparing the predicted mean RT data at the points of maximum and minimum conflict (Figure 3F) we would expect to see a clear difference (with RT data at the point of maximum conflict being slower), but the 95% HDI of the respective posterior predictives mostly overlap.

### Experiment 2: extension of the ‘pig’ dice game

Participants (N=50) performed a variant of the task in which the probability of losing at every die roll varied across conditions between 1/6, 2/6, and 3/6. As expected, the logistic regression analysis of choices made in this experiment revealed that (1) the cumulative sum of rewards was a good predictor for the probability of stopping in all conditions, as indicated by well-above-zero regression coefficients (HDI = [0.03, 0.05], [0.07, 0.09], [0.09, 0.14] for the 1/6, 2/6, and 3/6 conditions, respectively; Figure 4A), and (2) the average IP was substantially lower for the two added conditions (standard 1/6 condition: HDI = [108.75 189.74]; 2/6 condition: HDI = [56.83 92.89]; 3/6 condition: HDI = [26.56 49.21]; Figure 4B). The group-level IPs were all lower than the ‘risk-neutral’ IPs in their respective conditions, meaning that participants were on average risk-averse in all three conditions. Most importantly, the proportion of decisions made after the IP was substantially greater in the 2/6 condition compared to the 1/6 condition (with a mean difference equal to .17, p < .01, Tukey’s Test adjusted), and in the 3/6 condition compared to the standard 2/6 condition (with a mean difference equal to .31, p < .01, Tukey’s Test adjusted) (Figure 4C).

**Figure 4.**
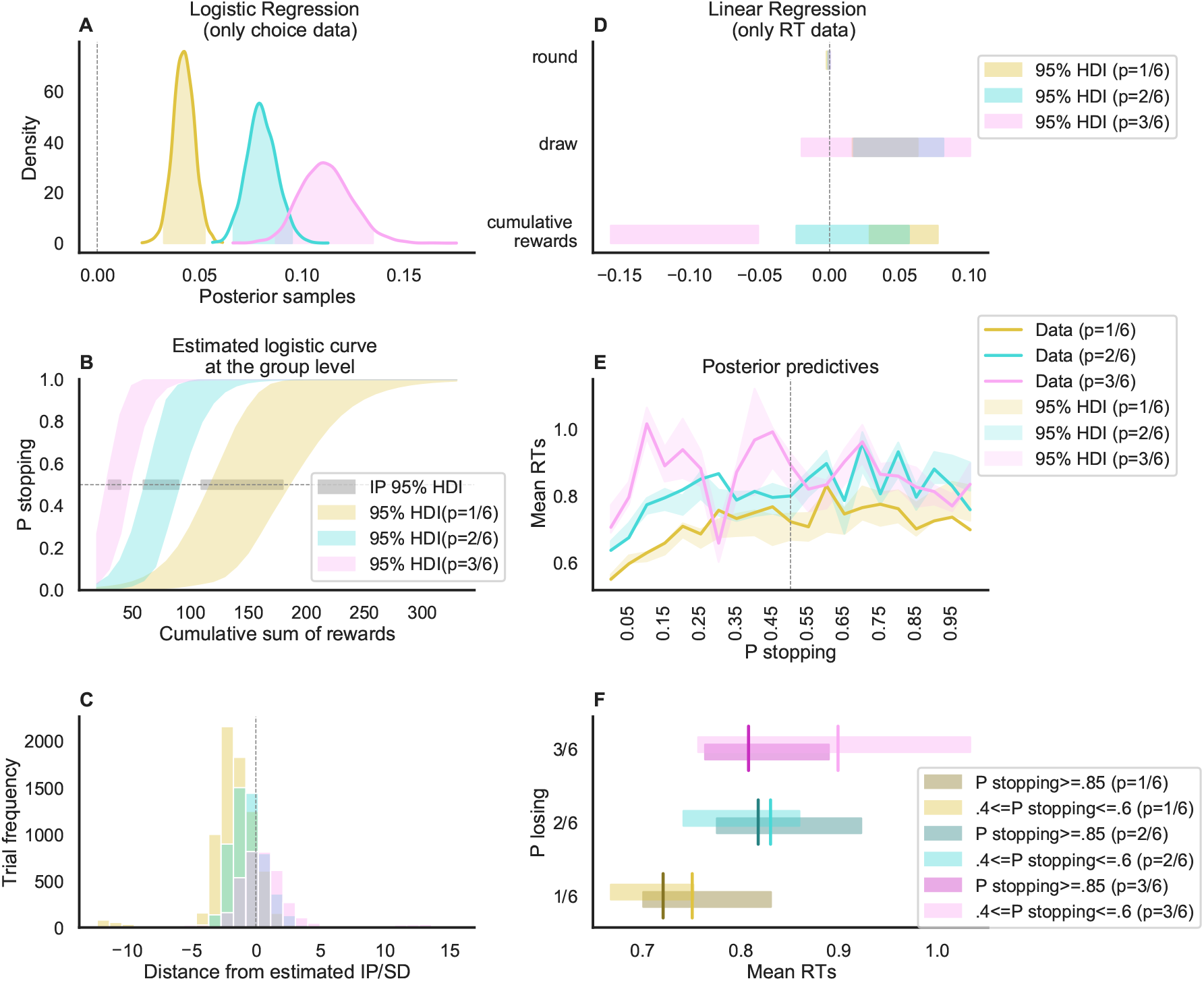
Regression analyses Experiment 2. The left column shows the results of the logistic model fit on choices, while the right column shows the results of the linear model fit on RTs. **A** Posterior distribution of the cumulative sum of rewards coefficients (separate per condition) at the group level (the shaded area is the 95% HDI), when we predict the probability of stopping within a round. **B** Estimated logistic curves (colored shaded area) and the IP (grey shaded area) at the group level. **C** Distribution of trials before and after the estimated IP, separately by condition. **D** Posterior distributions of the round number, draw number within a round, and cumulative sum of rewards coefficients at the group level (the shaded area is the 95% HDI), when we predict RTs.**E** Posterior predictives of mean RT as the probability of stopping increases within a round. **F** Comparison of the posterior predictives, selectively at the points of maximum conflict and at the points of maximum probability of stopping. Here the vertical lines represent the data while the shaded bars are the predictions.

As with Experiment 1, we tested the influence of round number, decision number, and cumulative sum of rewards on RT data (Table B2). Similar to Experiment 1, there was a negative effect of round number on RT in the 1/6 condition (implying faster decisions later in the experiment), but no systematic effects in the other two conditions. The coefficient for the decision number was above 0 for the 1/6 and 2/6 condition (implying slower decisions later in a trial). Most importantly, the cumulative sum of rewards coefficient was lower than 0 in the 3/6 condition, around 0 in the 2/6 condition, and positive in the 1/6 condition. This shows that the relationship between cumulative sum of rewards and RT critically depends on the task settings: As the proportion of decision after the IP increases, the effect of reward sum on RT flips from positive to negative.

We also looked at the posterior predictive distributions of RT as a function of the predicted probability to stop (Figure 4E and F). Here, we were specifically interested in comparing the predicted RT for a stop probability close to .5 (i.e., when being at the IP) against the predicted RT for a stop probability close to 1 (i.e., when being well beyond the IP). Even though on average the RT was lower at a stop probability close to 1 across all conditions, this analysis did not provide conclusive evidence for a decline of RT after the IP. Therefore, we proceeded with simultaneously fitting choice and RT data using a sequential sampling modeling approach.

### Sequential sampling modeling

The previous sections provide evidence that switch value and choice conflict are confounded in standard versions of the ‘pig’ dice game (and any games that share an analogous structure). While, under the appropriate experimental conditions (i.e., higher probabilities of loss), it is possible to increase the number of trials after the estimated IPs, it is still unclear whether this can help discriminate between monotonic or non-monotonic patterns in the RT data. This might be because we have thus far relied on separate regression analyses for choice and RT data. As a result, we did not take advantage of the joint information provided by both choice and RT data (and by the full shape of the RT distribution). At the same time, we also did not account for potential asymmetries in the speed with which one chooses to continue versus stop the round, and for ways in which different conditions may affect how cautious participants are in making their decisions. To address these gaps, we present and compare different variants of a process model for decisions made in this game, based on the sequential sampling modeling framework.

Sequential sampling models allow us to fit choice and RT data jointly in order to map them onto meaningful model variables and parameters. In essence, sequential sampling models assume that a decision emerges from a noisy process of evidence accumulation that is terminated as soon as a targeted level of evidence in favor of a particular choice option has been reached (Busemeyer, Gluth, Rieskamp, & Turner, 2019; Gold & Shadlen, 2007; Heekeren, Marrett, & Ungerleider, 2008; Ratcliff, Smith, Brown, & McKoon, 2016). We used the core version of the DDM (Ratcliff, 1978; Ratcliff & Rouder, 1998) with its four parameters drift rate, boundary separation, starting point, and non-decision time, but no across-trial variability parameters. The drift rate was linked to the utility difference between choosing to continue (vs. to stop) by specifying it at every decision point *t* as:

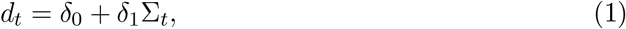

where ∑*_t_* refers to the cumulative sum of rewards, and *δ*_0_ and *δ*_1_ are free parameters. A positive drift rate favors the decision to stop, and a negative drift rate favors the decision to continue. It follows that this specification of the drift rate implicitly models the relationship between subject-specific levels of risk preferences and their IPs: As long as *δ*_0_ < *δ*_1_∑_*t*_, there is more evidence for continuing, but as soon as *δ*_0_ > *δ*_1_∑_*t*_, there is more evidence for stopping. At the IP, we have *δ*_0_ = *δ*_1_∑_*t*_, which implies that the probability to choose either option is .5, the expected RT is the longest, and choice conflict is highest. Therefore, we can use the DDM to infer participants’ IPs on the basis of their joint choice and RT data and model risk preferences similarly to how it is done in the logistic regression model. In addition to modeling the influence of accumulated reward on drift rate, we also generated variants of the DDM that allowed for the possibility that either the accumulated reward or the decision number might affect the threshold for responding (e.g., participants could become increasingly cautious after each ‘continue’ decision) or bias the starting point (e.g., the evidence required for making a ‘stop’ decision could become less and less after each ‘continue’ response).

The DDM variants were fitted to the choice and RT data of Experiment 2 using a hierarchical Bayesian modeling approach. Importantly, all models assumed different sets of parameters across the different conditions of the ‘pig’ dice game. In total, we compared 10 different DDM variants against each other, including a baseline model that assumed a fixed drift rate for every decision. As can be seen in Table 1, the DDM variant that provided the most parsimonious account of the data assumed that cumulative sum of rewards but not decision number impacted all of the considered DDM parameters (i.e., drift rate, threshold, and starting-point bias). Closer inspection of the posterior predictive distributions of the parameters coefficients (see Table B3 and Figure 5A) indicated that accumulated reward had the expected influence on the drift rate, with parameter being well above 0 in all three conditions.

**Table 1.** Model comparison of diffusion decision models (DDM) based on WAIC (data: Experiment 2).

| ID | Drift-rate | Starting-point | Threshold | $p_{\text{WAIC}}$ | -lppd | WAIC | WAIC<sub>se</sub> |
| --- | --- | --- | --- | --- | --- | --- | --- |
| 1 | not modulated | not modulated | not modulated | 486 | 6552 | 14076 | 273 |
| 2 | cumulative sum | not modulated | not modulated | 615 | 2903 | 7036 | 286 |
| 3 | cumulative sum | not modulated | decision number | 643 | 2821 | 6929 | 287 |
| 4 | cumulative sum | not modulated | cumulative sum | 652 | 2775 | 6854 | 288 |
| 5 | cumulative sum | decision number | not modulated | 646 | 2831 | 6955 | 286 |
| 6 | cumulative sum | decision number | decision number | 662 | 2776 | 6877 | 287 |
| 7 | cumulative sum | decision number | cumulative sum | 678 | 2713 | 6782* | 288 |
| 8 | cumulative sum | cumulative sum | not modulated | 652 | 2705 | 6714* | 290 |
| 9 | cumulative sum | cumulative sum | decision number | 683 | 2625 | 6616* | 291 |
| 10 | cumulative sum | cumulative sum | cumulative sum | 688 | 2601 | 6577** | 292 |

*Note.* If a DDM parameter was not modulated by any variable, a single value was estimated for it. If it was modulated, an intercept and a coefficient variable were estimated per variable. Lower WAICs indicate better fits to data after accounting for model complexity. \*\*Best model. \*These models do not fit credibly worse than the best model (because of the relatively high WAIC<sub>se</sub>).

**Figure 5.**
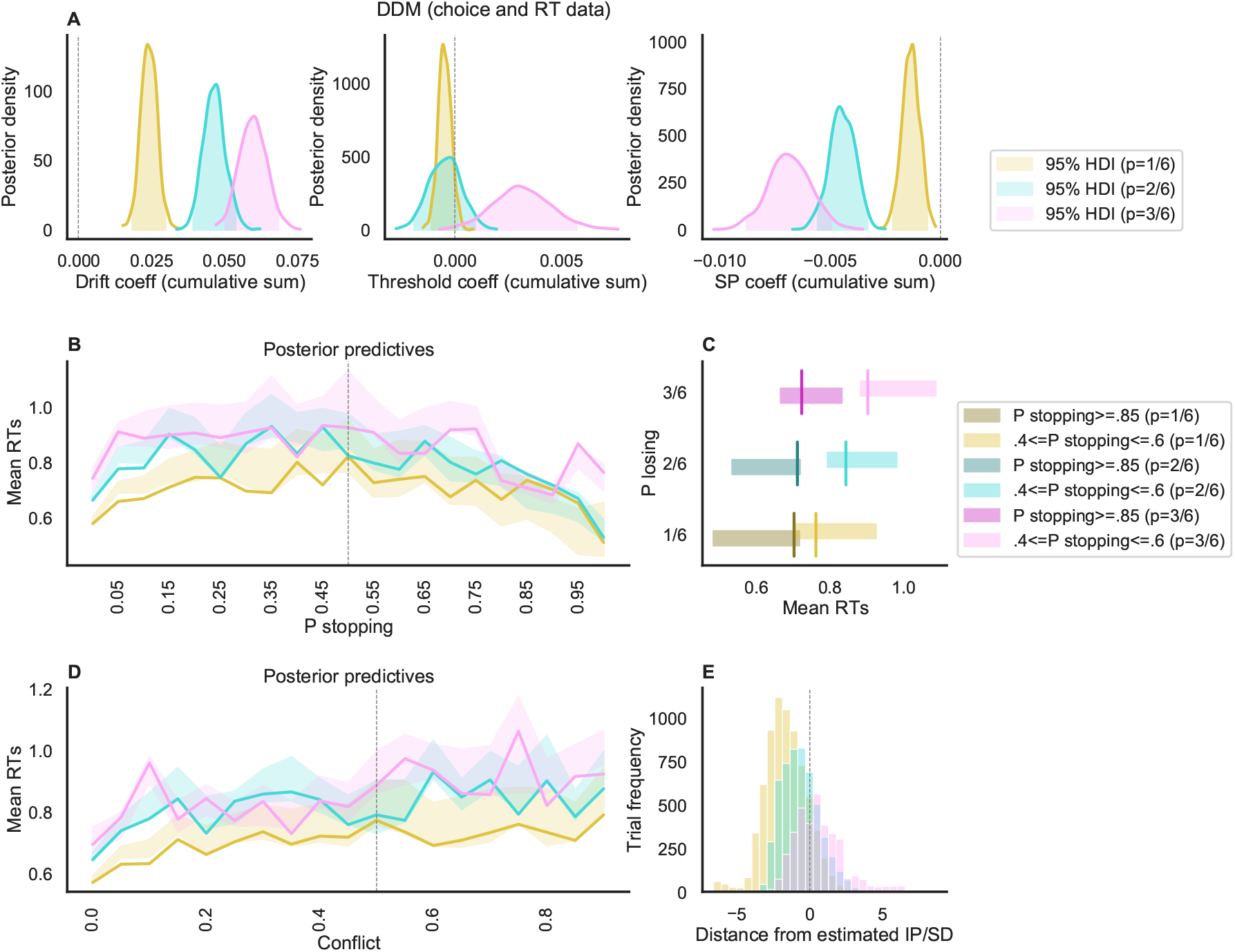
DDM analyses of Experiment 2. **A** Posterior distribution of the cumulative sum of rewards coefficients (separate per condition) at the group level (the shaded area is the 95% HDI) for three of the DDM parameters. **B** Posterior predictives of mean RT data as the probability of stopping increases within a round. **C** Comparison of the same posterior predictives, selectively at the points of maximum conflict and at the points of maximum probability of stopping. Here, the vertical lines represent the data while the shaded bars are the predictions. **D** Posterior predictives of mean RT data as conflict increases. **E** Distribution of trials after the estimated IP, separately by condition.

In the 3/6 condition but not the other two conditions, increases in the cumulative sum of rewards were also associated with higher thresholds, indicating greater overall caution when a lot of rewards had been collected. Across all three conditions, we also found that greater cumulative rewards were associated with a shift in one’s starting point towards the boundary for continuing (vs. stopping). While this may seem counter-intuitive at first (given that participants will also be increasingly likely to stop as cumulative reward increases), we speculate that it may reflect a bias towards repeating the previous response(s) (i.e., perseveration; Kool, Cushman, and Gershman (2016); Miller, Ludvig, Pezzulo, and Shenhav (2018); Miller, Shenhav, and Ludvig (2019)).

Analogous to the logistic regression analysis, we estimated the IP based on the drift-rate coefficients at a group-level and, based on them, the number of trials before and after the IP:

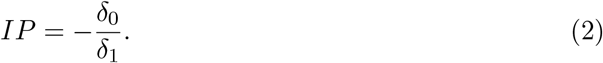

The estimated IP were similar to the ones obtained from the logistic regression: HDI = [115.00 190.98] (standard 1/6 condition); HDI = [61.67 89.19] (2/6 condition); and HDI = [27.50 45.91] (3/6 condition), confirming that participants were on average risk-averse in all three conditions. Accordingly, we also obtained a similar percentage of decisions made after the IP (Figure 5E): again, they were substantially more in the 2/6 condition compared to the 1/6 condition (with a mean difference equal to .14, p < .05, Tukey’s Test adjusted), and in the 3/6 condition compared to the standard 2/6 condition (with a mean difference equal to .32, p < .01, Tukey’s Test adjusted). For a direct comparison of the logistic and DDM coefficients, see the Appendix C).

In contrast to the linear regression results, our inspection of the posterior predictive distributions of RT as a function of the predicted probability to stop and of conflict (Figure 5B and D) allowed us to confirm one of our central hypotheses: The RT decreases after the IP, and thus reflect choice conflict rather than switch value. More specifically, when comparing the predicted RT for a stop probability close to .5 (i.e., when being at the IP) against the predicted RT for a stop probability close to 1 (i.e., when being well beyond the IP) the difference between mean RT is higher in our proposed conditions (i.e., conditions 2/6 and 3/6) compared to the original one (i.e., condition 1/6) (Figure 5C).

Taken together, the sequential sampling modeling analyses confirm that both choice and RT data in the ‘pig’ dice game are primarily driven by choice conflict, and that RT data exhibit a non-monotonic dynamic insofar as they first increase up to the IP but then decrease after it. Notably, this specific influence of choice conflict calls into question the conclusion of Meder and colleagues (2016), who concluded that RT (and with it, dACC activity) reflects switch value alone, but only on the basis of using the standard version of the dice task and analysing the data with regression analyses only. In addition (and more in line with Meder and colleagues), we also found evidence that the cumulative sum of rewards (and thus switch value) exerted multiple influences on behavior by affecting the drift rate, the starting-point bias, and (in the 3/6 condition) the boundary separation.

## Discussion

In this study, we sought to illustrate difficulties inherent to studies of sequential decision-making under risk, in teasing apart how people process the value of switching (stopping a given round and cashing out) from conflict over whether to stay or switch. We further demonstrated how these challenges can be overcome through a combination of cognitive modeling, model-driven experimental modifications, and advanced statistical analyses. To this end, we took the ‘pig’ dice game as an exemplary paradigm of sequential decision making. In this task, a player can pile up rewards over a sequence of repeated die rolls while facing a constant threat of losing all of the collected rewards. By analyzing the expected behavior of a rational (i.e., risk-neutral) agent, we demonstrated that the standard version of the task may not be able to disambiguate choice conflict and switch value (here conceptualized as the sum of collected rewards within a round) because of the limited number of decisions made after the IP. Thus, we extended the ‘pig’ dice game by adding two new conditions, in which we increased the probability of losing the cumulative sum of rewards. This modification led to a more balanced design of the task. Ultimately, the simultaneous modeling of choice and RT data with the DDM allowed us to identify both influences of choice conflict as well as switch value, and to map those influences onto different DDM parameters.

Our findings have broad implications for research into sequential decision-making in general and foraging-like settings in particular. With respect to research on foraging, our results confirm that the pig dice task, as originally devised, cannot disentangle contributions of switch value (cf. foraging value) from those of choice conflict when investigating underlying mechanisms, as Meder et al. (2016) attempted to do. While we did not collect neural data in this study, we did replicate their behavioral findings with an identical task design (Experiment 1) and showed that proper estimates of choice conflict for such a design are highly collinear with estimates of switch value. Our results are silent on which of these variables better accounts for dACC activity during such a task, but our normative and empirical findings from Experiment 2 suggest a clear path towards resolving this question. Previous studies by Shenhav and colleagues applied this same approach to deconfounding foraging value and choice conflict in a non-sequential foraging choice task (originating in Kolling et al. (2016)) and demonstrated that choice conflict consistently accounted for dACC activity during these choices better than foraging value (Shenhav, Straccia, et al., 2016; Shenhav et al., 2014; Zacharopoulos, Shenhav, Constantino, Maio, & Linden, 2018). It is therefore possible that a neuroimaging study of a deconfounded version of the pig dice task (e.g., our Experiment 2) would similarly demonstrate that activity previously attributed to switch value (e.g., cumulative sum of rewards) is better accounted for by choice conflict, but this awaits empirical testing. Importantly, in extending these theoretical and empirical findings to the domain of sequential choices, our current work provides an even richer account of relevant real-world choice settings, including how trial history effects within such settings help shape choice behavior over the course of a patch/round (see also Kane et al. (2021)).

There is a plethora of work showing that people can adapt their choice strategies to different environments (Gluth, Rieskamp, & Büchel, 2014; Payne, Bettman, & Johnson, 1988; Rieskamp & Otto, 2006; Todd & Gigerenzer, 2007). Thus, it is possible that participants used different strategies in the three different conditions. Take the 3/6 condition as an example, in which the loss probability at every decision point is 50%. Effectively, this eliminates the sequential element of the task, which is usually characterized by a sequence of multiple consecutive actions that are terminated by a final choice (see Introduction). Therefore, participants may have employed a choice strategy in this condition that is typical of and appropriate for standard 2-alternative-forced-choice tasks and that is well-captured by the DDM. In the other two conditions, however, they may have used a different strategy that could have resulted in different dynamics. Although we cannot rule out this possibility, our modeling results speak against the notion of different strategies, as the DDM provided a good account of the behavioral data across all conditions, and the choice dynamics remained stable in most aspects. Furthermore, we suppose that the within-subject design of Experiment 2, in which each participant was required to perform all three conditions, should have discouraged the adoption of different choice strategies from one condition to the next.

As stated above, the present work has strong connections to both the foraging literature and sequential decision making under risk and uncertainty. The latter does not only include the BART and the ‘pig’ dice game, but also the Angling Risk Task (Pleskac, 2008), the Columbia Card Task (Figner, Mackinlay, Wilkening, & Weber, 2009), and the stock-buying task used by Gluth, Rieskamp, and Büchel (2012). Such tasks have been used to study neural and physiological signals underlying evaluations of risky choice (see also Gluth, Rieskamp, and Büchel (2013); van Duijvenvoorde et al. (2015)). The current findings have very broad implications for research across these and other sequential decisionmaking tasks. First, the basic structure of sequential decision-making paradigms makes it difficult to investigate different dynamics, because most task features (e.g., cumulative rewards, escalating risks) and internal states (e.g., choice conflict, switch values) develop in a monotonic manner up to the terminal action. Therefore, it is necessary to understand this limitation and to consider variations of tasks that allow the researcher to investigate what happens after the point of maximum conflict has been reached. To exemplify how to apply our insights to other paradigms, take the BART: This task could be extended by a variant, in which a single pump occasionally increases the balloon’s volume massively. If this happens, the IP has most likely been exceeded to a large extent, and the decision to switch to the next round is fairly easy and made with comparatively little choice conflict.

Our results also demonstrate the utility of combining state-of-the-art statistical analyses of behavior (e.g., hierarchical Bayesian techniques) with similarly sophisticated cognitive modeling approaches in order to maximize the potential of dissociating different dynamics. For instance, our DDM results suggest that switch value not only influences the rate of evidence accumulation within a given trial but also the ‘initial settings’ of the decision process in terms of boundary separation and starting-point bias. In particular, our modeling results suggest the presence of a perseveration (or stickiness) effect of increasing trials within a given round, which leads to fast errors (i.e., fast “continue” choices late in the round, in the presence of high evidence for “stopping”). This was made evident by a modulation of the starting point bias by the cumulative rewards in each round. This effect would have not been evident with simple regression analyses, which do not take the whole distribution into account. It will be important for future research to try to generalize these results to other sequential decision-making scenarios and to identify the (potentially diverging) neural correlates of different cognitive mechanisms.

## Methods

### Simulations and task extension

We performed the first simulation to show how a risk-neutral, EV maximizing participant would behave in our extended version of the ‘pig’ dice game. Because we also simulated players with non-neutral risk preferences (see further below), we will use the more general term of expected utility (EU) in the following. The EU of the two options (i.e., stop vs. continue) were calculated as:

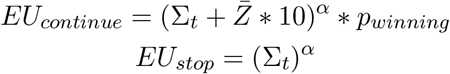

where 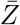 is the average winning number in a specific condition. In the 1/6 condition, we have 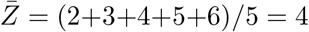; in the 2/6 condition, we have 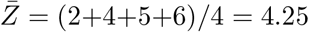; in the 3/6 condition, we have 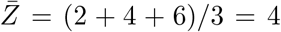. Parameter *α* was set to 1 for the risk-neutral player, so that the EU was equal to EV.

We simulated choices drawing random samples from a Bernoulli distribution, where the probability to stop was obtained using the softmax function:

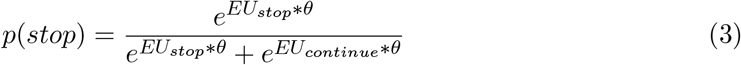

where *θ* is a sensitivity parameter, and was set to 0.2. We simulated 120 minutes of game play per condition.

The IPs of a risk-neutral player for each condition can be obtained by equating *EU_continue_* and *EU_stop_*:

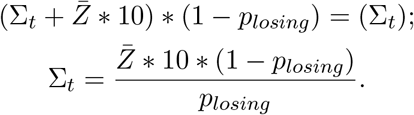

Therefore, when the probability of losing is 1/6:

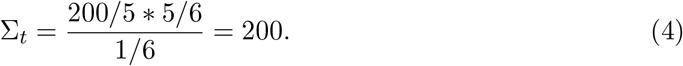

Likewise, when the probability of losing is 2/6:

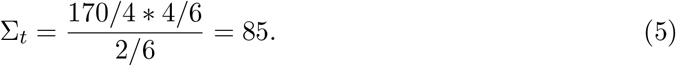

Finally, when the probability of losing is 3/6:

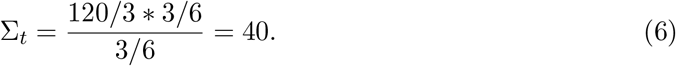

We performed the second simulation to show how a group of participants with different risk preferences would behave in the standard and extended versions of the ‘pig’ dice game. Instead of fixing parameter *α* at 1, we randomly drew 100 *α* values from a transformed normal distribution *α* ~ log(1 + exp(*N*(*M* = 0.5, *SD* = 0.3)), so that the resulting *α* had a mean of 1, a minimum value of 0.6 and a maximum value of 1.4. This distribution ensured that on average participants were risk neutral and had an IP around 200 when the probability of losing was 1/6. As for the risk-neutral player, the sensitivity *θ* was set to .2. However, to make the simulations more similar to the original task, we simulated 20 minutes of game play per participant and condition.

### Experiment 1

#### Participants

In Experiment 1, we tested 30 participants (22 females, age: 18-48, M=26.1, SD=6.9). Four participants were excluded, because they did not understand the task (e.g., they made less than 5% either continue or stop decision, or they made more “stop” than “continue” responses at the beginning of a round). The sample size was selected based on previous work with comparable sample sizes, but was not determined by a formal power analysis. Notably, the study that we sought to replicate (Meder et al., 2016) tested only 20 participants and analyzed the data of only 18 participants. Participants provided written informed consent and the study was approved by the Department of Psychology Ethics Committee at the University of Basel.

#### Experimental procedures

After reading and signing the consent form, participants were instructed on the ‘pig’ dice game. This task was played in multiple rounds for a predetermined amount of time. Following Meder and colleagues, participants played the task for 25 min. Each round consisted of multiple sequential decisions to continue rolling a regular six-sided die (and thereby accumulating rewards) or to cash in the cumulative sum of rewards by stopping the current round and proceeding to a new round. The two buttons to “continue” and to “stop” were counterbalanced across participants. As long as any number between 2 and 6 was rolled, the rolled number was multiplied by 10, and the reward was added to the current round’s cumulative sum of rewards. As soon as a 1 was rolled, all cumulative sum of rewards were lost, and the player automatically proceeded to a new round. Participants were informed that they would receive a bonus payment on top of their show-up fee (of 20 Swiss Francs per hour or course credits), and that the amount of this bonus was a function of the average cumulative sum of rewards per round, including the rounds in which all rewards were lost. More specifically, participants were paid out a tenth of this reward average as Swiss Francs. This incentive structure implies a simple and straightforward definition of the optimal policy for an EV maximizing, risk-neutral player, which is to attempt to accumulate a reward of 200 and then to cash in (see Methods above).

Each round began with a first die roll that was played out automatically. A regular white die with black dots was presented on a green, circular piece of carpet surrounded by a wooden background (Figure 2A). The roll of the die was animated by switching its orientation (tilted by 9 degrees to the right and to the left of the vertical axis) and randomly changing its displayed dots every 150 ms for a variable amount of time, drawn from a uniform distribution between 1.5 and 3.75 s and discretized in steps of 150 ms. Afterwards, the outcome of the current roll was displayed for 2 s by presenting the die in regular orientation. At the same time, the cumulative sum of rewards of the current round was presented on top of the die. In case of rolling a number from 2 to 6, participants had to make their decision to continue or to stop within this 2 s time window. Otherwise, they would be shown the losing die with a 0 reward on top for 2 seconds. In case the participant decided to stop, the cumulative sum of rewards of the current round was shown in the middle of the die for 2.5 s, and a new round started. If the participant decided to continue, the next die roll was animated as described above. After 25 min, the experiment ended (even if a round was not completed). Participants were then shown their average cumulative sum of rewards per round together with the respective bonus payments, received this payment together with their show-up fee, were debriefed, and left the experiment. The task was programmed using Psychopy (Peirce, 2007).

#### Statistical analyses

To estimate the IP for each participant, we performed a hierarchical Bayesian logistic regression analysis that regressed the decision to continue (vs. to stop) onto the cumulative sum of rewards (∑_*t*_) at every decision *t*. The regression model has the form:

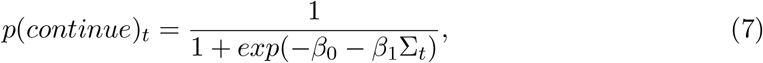

with *β*_0_ and *β*_1_ representing the intercept and slope coefficients, respectively. The IP, which is defined as the amount of cumulative sum of rewards at which *p*(*continue*)*_t_* = .5, can then be derived as:

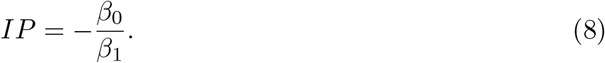

We adopted a hierarchical Bayesian approach to estimate the regression coefficients and the IP. Individual coefficients were drawn from group-level Normal distributions:

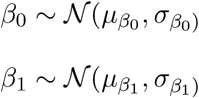

Mean and standard deviations of these distributions were themselves drawn using the following hyper-priors:

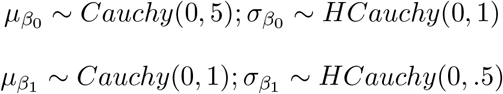

To investigate how RT of stop and continue decisions were affected by different dynamics including choice conflict and switch value, we performed a hierarchical Bayesian linear regression predicting the log(RT) by the variables round number (i.e., *r*), within-round decision number, cumulative sum of rewards (i.e., ∑_*t*_), and choice conflict (i.e., *cc_t_*). Choice conflict was defined as the derivative of the logistic curve taken from the aforementioned analysis on choice data (note that the slope/derivative of this curve is maximal at *p*(*stop*)*_t_* = .5 and falls of symmetrically for both smaller and greater values of *p*(*stop*)*_t_*). Thus, the linear regression of RT had the form:

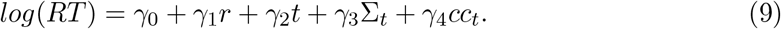

Individual regression coefficients were drawn from group-level normal distributions:

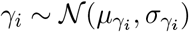

with hyper-priors for the coefficients *i* > 0:

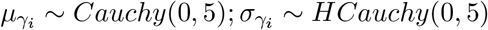

and for *i* = 0 (the intercept):

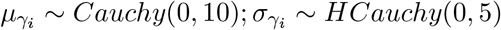

All Bayesian models were fit using PyStan 2.19, a Python interface to Stan (Carpenter et al., 2017). We ran two parallel chains until convergence was reached (1000 iterations per chain after 1000 burnin period). To assess convergence, we checked that the Gelman-Rubin convergence diagnostic 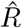 (Gelman & Rubin, 1992) was lower or equal to 1.01.

### Experiment 2

#### Participants

In Experiment 2, we tested 50 participants (35 females, age: 19-36, M=22.8, SD=3.2). Nine participants were excluded, using the same criteria as in Experiment 1. A larger sample size than in Experiment 1 was chosen because of the additional 2 conditions of the task and to counteract the problem of the low number of decisions after the IP per participant. However, the sample size was not determined by a formal power analysis. As with Experiment 1, participants provided written informed consent and the study was approved by the Department of Psychology Ethics Committee at the University of Basel.

#### Experimental procedures

The experimental procedures were largely identical to those of Experiment 1. The only exception was that the task included three different conditions, which varied the probability of losing the cumulative sum of rewards at every decision. Thus, in addition to the standard (1/6) condition, in which all rewards in a certain round were lost when rolling a 1, there was also a 2/6 condition, in which all rewards were lost in a round when rolling a 1 or a 3, and a 3/6 condition, in which all rewards in a round were lost when rolling any odd number (i.e., 1, 3, 5). Moreover, participants played 18 minutes of each condition, so that the entire experiment would not be too long. The order of conditions was counterbalanced across participants. In particular, we had three possible orders: high to low probability (p=3/6, p=2/6, p=1/6), low to high probability (p=1/6, p=2/6, p=3/6), and middle then high then low probability (p=2/6, p=3/6, p=1/6). We also counterbalanced across participants whether the “stop” button was on the right or on the left of the “continue” button. Participants were instructed about the presence of different conditions prior to the task.

#### Statistical analyses

Statistical analyses of Experiment 2 were largely identical to those of Experiment 1, with performing hierarchical Bayesian logistic and linear regressions for the choice and RT data, respectively. Note that separate regressors were estimated for each task condition.

#### Sequential sampling modeling

The DDM is arguably the most prominent representative of sequential sampling models. Without assuming across-trial variability in parameters (Ratcliff & Rouder, 1998), the DDM maps choice and RT data onto the four parameters drift rate *d_t_*, boundary separation *a*, non-decision time *T_er_*, and (relative) starting point *b* on the basis of the Wiener distribution (Navarro & Fuss, 2009):

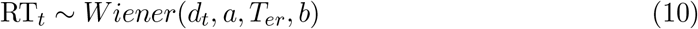

with:

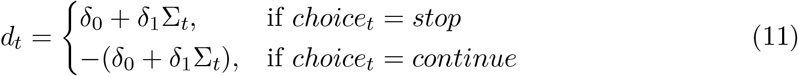

DDM parameters were estimated using hierarchical Bayesian modeling procedures (similarly to the logistic and linear regression models described above). Individual coefficients for drift rate, threshold and starting-point bias were drawn from group-level Normal distributions:

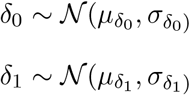

Mean and standard deviations of these distributions for drift rate and threhsold were themselves drawn using the following hyper-priors:

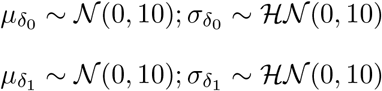

The hyper-priors of the starting-point bias were, however set to:

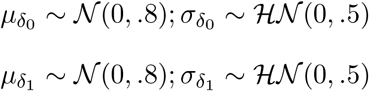

Since the non decision time did not have coefficients, its prior and hyper priors were the same as the *δ*_0_ parameter of drift rate and threshold parameters.

The DDM was applied to the data of Experiment 2, assuming different parameter sets for each condition.

#### Data and code availability

Data of all participants included in the final samples of the two experiments and custom code that supports the findings of this study is available at osf.io/meurt/.

## Appendix A Simulations with risk averse participants

**Figure A1.**
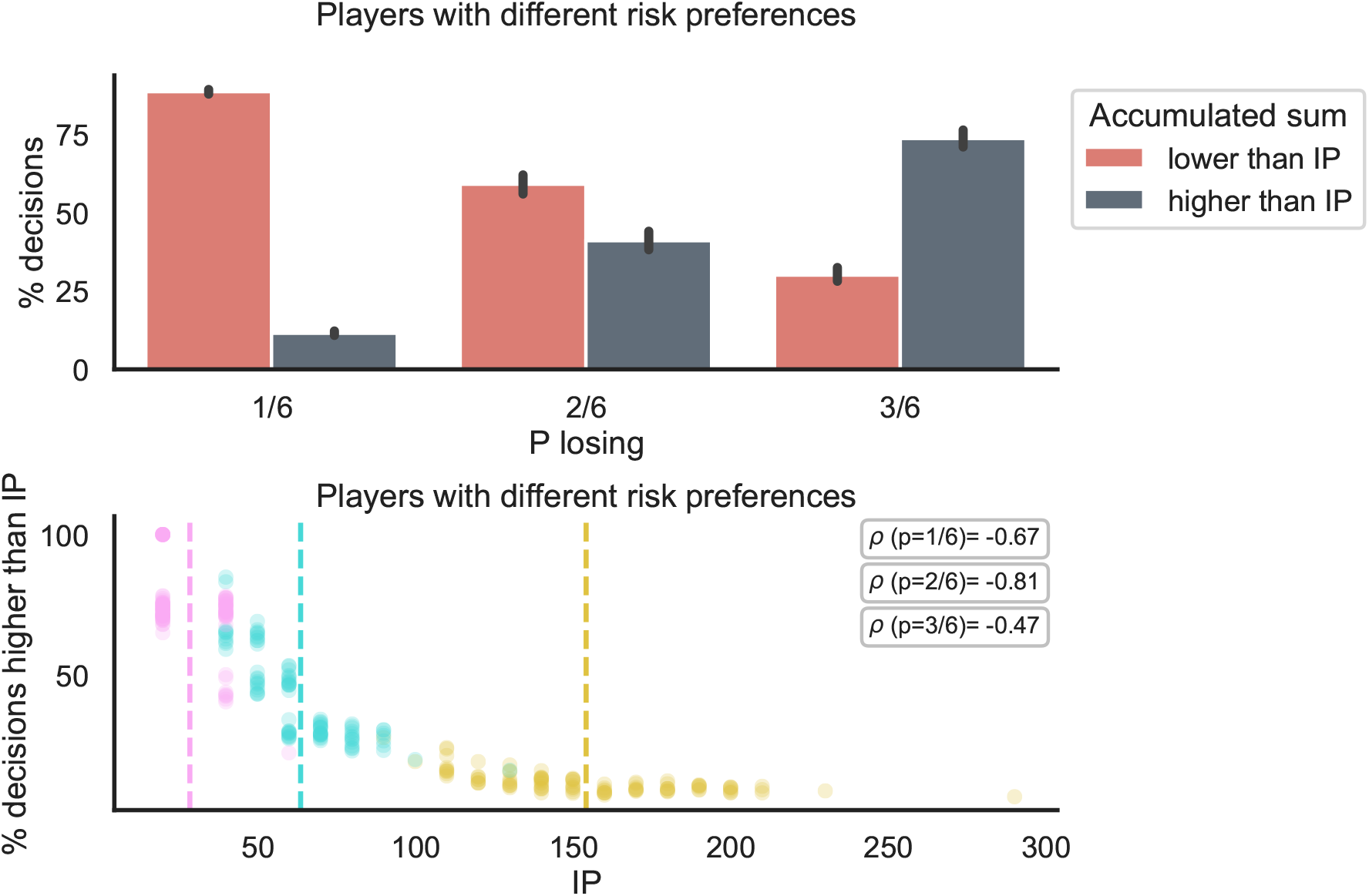
**Top panel:** Percentage of decisions before and after the IP for a simulated group of 100 players with different risk preferences (risk averse on average, with *α*=.8). The decisions before the IP significantly decrease with the probability of losing, making the task more balanced. **Lower panel**: Percentage of decisions after the IP as a function of the IP, for the same group of simulated participants as in C. Participants with higher IPs (more risk seeking) experience a more imbalanced task in the 2/6 and 1/6 conditions.

## Appendix B Regression coefficients

**Table B1.** Logistic regression coefficients summary.

| Experiment | Condition | Coefficient | $\mu$ HDI | $\sigma$ HDI |
| --- | --- | --- | --- | --- |
| 1 | 1/6 | intercept | [-5.7 -3.87] | [1.52 2.95] |
| 1 | 1/6 | cumulative sum | [0.03 0.05] | [0.02 0.03] |
| 2 | 1/6 | intercept | [-6.9 -5.4] | [1.68 2.99] |
| 2 | 1/6 | cumulative sum | [0.03 0.05] | [0.02 0.04] |
| 2 | 2/6 | intercept | [-6.97 -5.01] | [2.27 3.88] |
| 2 | 2/6 | cumulative sum | [0.06 0.09] | [0.03 0.06] |
| 2 | 3/6 | intercept | [-4.89 -2.92] | [2.26 3.91] |
| 2 | 3/6 | cumulative sum | [0.09 0.14] | [0.05 0.1] |

*Note.* Summary of the intercept and cumulative sum coefficients at the group level.

**Table B2.** Linear regression coefficients summary.

| Experiment | Condition | Coefficient | $\mu$ HDI | $\sigma$ HDI |
| --- | --- | --- | --- | --- |
| 1 | 1/6 | intercept | [-0.34 -0.21] | [0.14 0.24] |
| 1 | 1/6 | cumulative sum | [-0.004 0.049] | [0.035 0.075] |
| 1 | 1/6 | decision number | [0.02 0.07] | [0.02 0.06] |
| 1 | 1/6 | round number | [-0.070 -0.001] | [0.067 0.117] |
| 2 | 1/6 | intercept | [-0.52 -0.27] | [0.24 0.48] |
| 2 | 1/6 | cumulative sum | [0.03 0.07] | [0.04 0.08] |
| 2 | 1/6 | decision number | [0.02 0.06] | [0.03 0.07] |
| 2 | 1/6 | round number | [-0.0021 -0.0006] | [0.002 0.003] |
| 2 | 2/6 | intercept | [-0.38 -0.16] | [0.26 0.46] |
| 2 | 2/6 | cumulative sum | [-0.02 0.06] | [0.07 0.14] |
| 2 | 2/6 | decision number | [0.02 0.08] | [0.03 0.1] |
| 2 | 2/6 | round number | [-0.001 0.0001] | [0.001 0.002] |
| 2 | 3/6 | intercept | [-0.35 -0.17] | [0.19 0.35] |
| 2 | 3/6 | cumulative sum | [-0.16 -0.05] | [0.10 0.19] |
| 2 | 3/6 | decision number | [-0.02 0.10] | [0.11 0.22] |
| 2 | 3/6 | round number | [-0.0007 0.0004] | [0.001 0.002] |

*Note.* Summary of the intercept, cumulative sum of rewards, decision number, and round number coefficients at the group level.

**Table B3.** Diffusion decision model coefficients summary.

| Condition: | 1/6 | 2/6 | 3/6 |
| --- | --- | --- | --- |
| IP(low) | 115 | 61.67 | 27.5 |
| IP(high) | 190.98 | 89.19 | 45.91 |
| $\delta_1$(low) | 0.02 | 0.04 | 0.05 |
| $\delta_1$(high) | 0.03 | 0.05 | 0.07 |
| $\delta_0$(low) | -4.11 | -3.91 | -2.73 |
| $\delta_0$(high) | -3.33 | -3.03 | -1.74 |
| $A_1$(low) | -0.001 | -0.0018 | 0.0008 |
| $A_1$(high) | 0.0002 | 0.001 | 0.0061 |
| $A_0$(low) | 1.33 | 1.36 | 1.23 |
| $A_0$(high) | 1.57 | 1.61 | 1.47 |
| $z_1$(low) | -0.002 | -0.006 | -0.009 |
| $z_1$(high) | -0.001 | -0.003 | -0.005 |
| $z_0$(low) | 0.04 | 0.18 | 0.19 |
| $z_0$(high) | 0.22 | 0.35 | 0.37 |

*Note.* The IP was calculated based on the drift-rate coefficients alone ( $IP = -\delta_0/\delta_1$ ). The $\mu$ HDI are reported for all the diffusion decision model coefficients, that describe the effect of the cumulative sum on the main parameters: the drift-rate $\delta$ , the threshold $A$ , and the relative starting point $z$ . The model was only fit to the data of Experiment 2.

## Appendix C Qualitative comparison between the logistic and the sequential sampling modeling results

**Figure C1.**
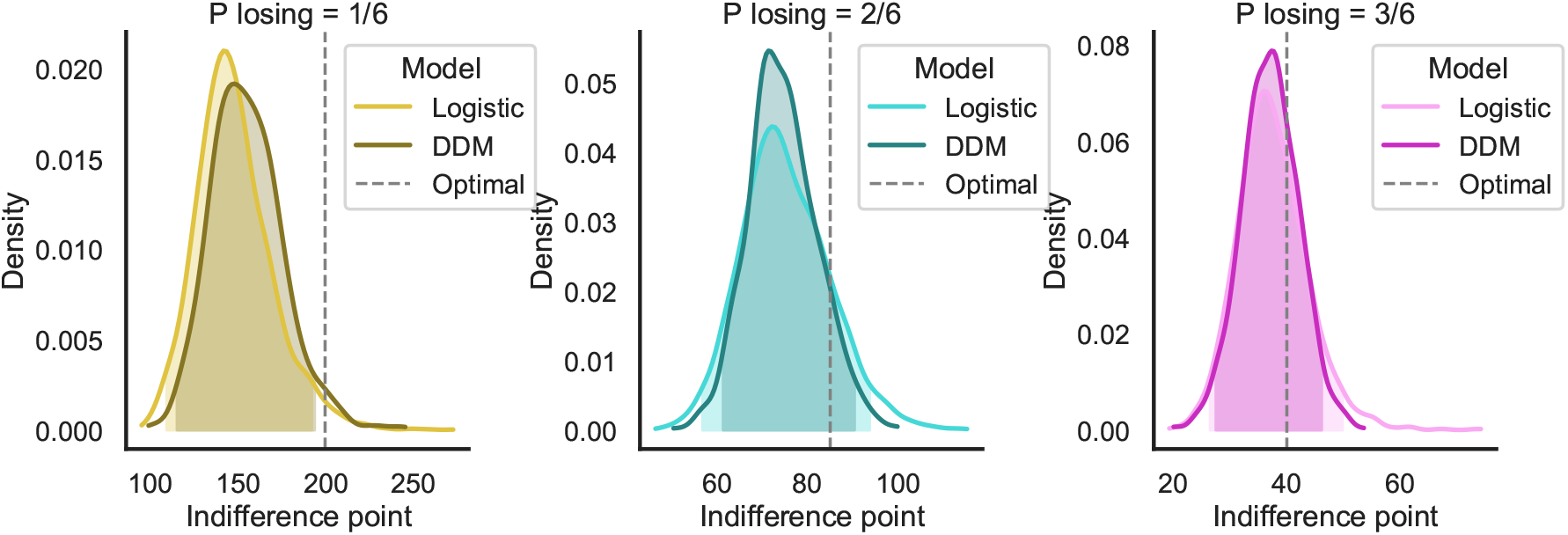
Posterior distribution of the group-level IP according to the logistic model and to the best fitting diffusion decision model in the second Experiment, separately for the different conditions. Note that the IP in the diffusion model is calculated based on the drift rate coefficients only.

## Notes

https://osf.io/meurt/

### Competing Interest Statement

The authors have declared no competing interest.

### Summary of Updates

I have changed the author list to match the one in the manuscript.

## References

Busemeyer, J. R., Gluth, S., Rieskamp, J., & Turner, B. M. (2019). Cognitive and neural bases of multi-attribute, multi-alternative, value-based decisions. Trends in Cognitive Sciences, 23, 251–263.

Carpenter, B., Gelman, A., Hoffman, M. D., Lee, D., Goodrich, B., Betancourt, M., … Riddell, A. (2017). Stan: A probabilistic programming language. Journal of Statistical Software, 76(1), 1–32. doi: 10.18637/jss.v076.i01

Charnov, E. L. (1976). Optimal foraging, the marginal value theorem. Theoretical Population Biology, 9, 129–136.

Figner, B., Mackinlay, R. J., Wilkening, F., & Weber, E. U. (2009). Affective and deliberative processes in risky choice: age differences in risk taking in the Columbia Card Task. Journal of Experimental Psychology. Learning, Memory, and Cognition, 35, 709–730.

Gelman, A., & Rubin, D. B. (1992). Inference from iterative simulation using multiple sequences. Statistical Science, 7(4), 457–472. doi: 10.1214/ss/1177011136

Gluth, S., Rieskamp, J., & Büchel, C. (2012). Deciding when to decide: time-variant sequential sampling models explain the emergence of value-based decisions in the human brain. Journal of Neuroscience, 32, 10686–10698.

Gluth, S., Rieskamp, J., & Büchel, C. (2013). Classic EEG motor potentials track the emergence of value-based decisions. NeuroImage, 79, 394–403.

Gluth, S., Rieskamp, J., & Büchel, C. (2014). Neural evidence for adaptive strategy selection in value-based decision-making. Cerebral Cortex, 24, 2009–2021.

Gold, J. I., & Shadlen, M. N. (2007). The neural basis of decision making. Annual Review of Neuroscience, 30, 535–574.

Heekeren, H. R., Marrett, S., & Ungerleider, L. G. (2008). The neural systems that mediate human perceptual decision making. Nature Reviews Neuroscience, 9, 467–479.

Kane, G. A., James, M. H., Shenhav, A., Daw, N. D., Cohen, J. D., & Aston-Jones, G. (2021). Rat anterior cingulate cortex continuously signals decision variables in a patch foraging task. bioRxiv. Retrieved from https://www.biorxiv.org/content/early/2021/09/26/2021.06.07.447464 doi: 10.1101/2021.06.07.447464

Kolling, N., Behrens, T. E., Mars, R. B., & Rushworth, M. F. (2012). Neural mechanisms of foraging. Science, 336(6077), 95–98.

Kolling, N., Wittmann, M. K., Behrens, T. E. J., Boorman, E. D., Mars, R. B., & Rushworth, M. F. S. (2016). Value, search, persistence and model updating in anterior cingulate cortex. Nature Neuroscience, 19.

Kool, W., Cushman, F. A., & Gershman, S. J. (2016). When does model-based control pay off? PLoS computational biology, 12(8), e1005090.

Lejuez, C. W., Read, J. P., Kahler, C. W., Richards, J. B., Ramsey, S. E., Stuart, G. L., … Brown, R. A. (2002). Evaluation of a behavioral measure of risk taking: The Balloon Analogue Risk Task (BART). Journal of Experimental Psychology: Applied, 8, 75–84.

Meder, D., Haagensen, B. N., Hulme, O., Morville, T., Gelskov, S., Herz, D. M., … Siebner, H. R. (2016). Tuning the brake while raising the stake: network dynamics during sequential decision-making. Journal of Neuroscience, 36(19), 5417–5426.

Miller, K. J., Ludvig, E. A., Pezzulo, G., & Shenhav, A. (2018). Realigning models of habitual and goal-directed decision-making. In Goal-directed decision making (pp. 407–428). Elsevier.

Miller, K. J., Shenhav, A., & Ludvig, E. A. (2019). Habits without values. Psychological review, 126(2), 292.

Navarro, D. J., & Fuss, I. G. (2009). Fast and accurate calculations for first-passage times in Wiener diffusion models. Journal of Mathematical Psychology, 53, 222–230.

Payne, J. W., Bettman, J. R., & Johnson, E. J. (1988). Adaptive strategy selection in decision making. Journal of Experimental Psychology: Learning, Memory, and Cognition, 14, 534–552.

Peirce, J. W. (2007). Psychopy—psychophysics software in python. Journal of neuroscience methods, 162(1-2), 8–13.

Pleskac, T. J. (2008). Decision making and learning while taking sequential risks. Journal of Experimental Psychology: Learning, Memory, and Cognition, 34, 167–185.

Ratcliff, R. (1978). A theory of memory retrieval. Psychological Review, 85(2), 59–108. doi: 10.1037/0033-295X.85.2.59

Ratcliff, R., & Rouder, J. N. (1998). Modeling response times for two-choice decisions. Psychological Science, 9(5), 347–356. doi: 10.1111/1467-9280.00067

Ratcliff, R., Smith, P. L., Brown, S. D., & McKoon, G. (2016). Diffusion decision model: current issues and history. Trends in Cognitive Sciences, 20, 260–281.

Rieskamp, J., & Otto, P. E. (2006). SSL: a theory of how people learn to select strategies. Journal of Experimental Psychology: General, 135, 207–236.

Schonberg, T., Fox, C. R., Mumford, J. A., Congdon, E., Trepel, C., & Poldrack, R. A. (2012). Decreasing ventromedial prefrontal cortex activity during sequential risk-taking: An fMRI investigation of the balloon analog risk task. Frontiers in Neuroscience, 6.

Shenhav, A., Cohen, J. D., & Botvinick, M. M. (2016). Dorsal anterior cingulate cortex and the value of control. Nature neuroscience, 19(10), 1286–1291.

Shenhav, A., Straccia, M. A., Botvinick, M. M., & Cohen, J. D. (2016). Dorsal anterior cingulate and ventromedial prefrontal cortex have inverse roles in both foraging and economic choice. Cognitive, Affective, & Behavioral Neuroscience, 16(6), 1127–1139.

Shenhav, A., Straccia, M. A., Cohen, J. D., & Botvinick, M. M. (2014). Anterior cingulate engagement in a foraging context reflects choice difficulty, not foraging value. Nature Neuroscience, 17(9), 1249.

Todd, P. M., & Gigerenzer, G. (2007). Environments that make us smart: ecological rationality. Current Directions in Psychological Science, 16, 167–171.

van Duijvenvoorde, A. C. K., Huizenga, H. M., Somerville, L. H., Delgado, M. R., Powers, A., Weeda, W. D., … Figner, B. (2015). Neural correlates of expected risks and returns in risky choice across development. Journal of Neuroscience, 35, 1549–1560.

Zacharopoulos, G., Shenhav, A., Constantino, S., Maio, G. R., & Linden, D. E. (2018). The effect of self-focus on personal and social foraging behaviour. Social cognitive and affective neuroscience, 13(9), 967–975.

